# Dermal nerve growth factor is increased in prurigo nodularis compared to atopic dermatitis

**DOI:** 10.1101/2022.06.15.496275

**Authors:** Junwen Deng, Varsha Parthasarathy, Zachary Bordeaux, Melika Marani, Kevin Lee, Chi Trinh, Nishadh Sutaria, Hannah L. Cornman, Anusha Kambala, Thomas Pritchard, Shihua Chen, Olusola O. Oladipo, Madan M. Kwatra, Martin P. Alphonse, Shawn G. Kwatra

## Abstract

**Background:** Prurigo nodularis (PN) is a chronic, pruritic, inflammatory skin disease characterized by hyperkeratotic nodules on the trunk and extremities. While there is growing research on the immunological basis of PN, the neuropathic and structural components of PN lesions are unknown.

**Objective:** To determine the inflammatory, neuropathic, and structural pathways in PN compared to atopic dermatitis (AD).

**Methods:** Lesional and non-lesional skin biopsies were collected from 13 PN and 6 AD patients. mRNA and protein expression in biopsies was determined using RNA-Sequencing and immunohistochemistry (IHC), respectively. Differentially expressed genes (DEGs) were identified using the *DESeq2* R package and pathway level enrichment was determined using Gene Set Enrichment Analysis. IHC expression was quantified with QuPath followed by statistical comparison with the Student’s t-test and Mann-Whitney *U*.

**Results:** Compared to lesional AD, lesional PN had greater mRNA expression of MMPs, OSM, NGF, IL1β, CXCL2, CXCL5, CXCL8, and insulin-like growth factors, and lower expression of CCL13, CCL26, EPHB1, and collagens. Compared to non-lesional AD, non-lesional PN showed upregulation of keratin-family genes. GSEA revealed that lesional PN had greater keratinization, cornified envelope, myelin sheath, TGF-beta signaling, extracellular matrix disassembly, metalloendopeptidase activity, and neutrotrophin-TRK receptor signaling, while non-lesional PN had higher keratin filament, extracellular structure organization, extracellular matrix disassembly, and angiogenesis. IHC showed increased dermal nerve growth factor (NGF) expression in lesional PN compared to lesional AD (p=0.038), and greater epidermal NGF compared to dermal NGF in non-lesional PN (p=0.014).

**Limitations:** Single, tertiary care center.

**Conclusions:** PN demonstrated increased neurotrophic and extracellular matrix (ECM) remodeling signatures compared to AD, possibly explaining the morphological differences in their lesions. These signatures may therefore be important components of the PN pathogenesis and may serve as therapeutic targets.

---

Prurigo nodularis (PN) is a chronic, pruritic inflammatory skin disease characterized by hyperkeratotic nodules on the extensor surfaces and trunk.^1^ PN features both inflammatory and neuropathic dysregulation and shares several pathogenic features with atopic dermatitis (AD), including cutaneous upregulation of interleukin (IL)-4R and Th22 transcriptomic signatures.^1,2^ However, the exact pathogenesis of PN is not well described. In particular, the role of nerve growth factor (NGF), which regulates nerve development, has not been previously examined in PN in relation to AD. Therefore, we hypothesized that direct comparison of the cutaneous transcriptomes and immunohistochemical distribution of NGF in PN and AD patients would provide insight into the unique inflammatory and neuropathic mechanisms of PN.

This study was performed through transcriptomic and immunohistochemical analysis of skin punch biopsies from lesional and non-lesional areas of PN and AD patients with moderate-to-severe pruritus. Transcriptomic analysis was performed on a total of 38 lesional and non-lesional skin biopsies, including 13 PN (mean age 54.8±14.2 years, 84.6% female, and 76.9% African American) and 6 AD (mean age 55.2±15.8 years, 83.3% female, and 100.0% African American) patients. Full sequencing methodology can be found in our prior articles.^1,2^. Normalization and differentially expressed gene (DEG) calculations were conducted using *DESeq2* (Bioconductor). DEGs were defined as genes with a log2-fold change <-1.5 or >1.5. The false discovery rate (FDR) was calculated to control for multiple hypothesis testing. Pathway-level comparisons were performed using Gene Set Enrichment Analysis (GSEA).

Immunohistochemistry (IHC) staining for NGF was performed on formalin fixed, paraffin embedded sections on age-, sex-, and race-matched lesional PN and AD samples (n=8 each) and matched non-lesional samples (n=3 each). Epitope retrieval was performed using Ventana Ultra CC1 buffer (catalog# 6414575001, Roche). Anti NGF (1:500 dilution; catalog# ab52918, Abcam) primary antibody was applied and detected using an anti-rabbit HQ detection system (catalog# 7017936001 and 7017812001, Roche) followed by Chromomap DAB IHC detection kit (catalog# 5266645001, Roche) and counterstaining with Mayer’s hematoxylin. Quantitative analysis of the percentage of NGF-positive cells was performed using QuPath. Normality was analyzed using a Shapiro-Wilk test, and an unpaired T-test was performed for normally distributed data sets and a Mann-Whitney U test was performed for non-normal data sets.

Transcriptome analysis revealed 1,415 DEGs between lesional PN and AD skin (PN/AD L), 42 DEGs between non-lesional PN and AD skin (PN/AD NL), and 6 DEGs in common between PN/AD L and PN/AD NL (Fig. 1A-B). Comparing lesional PN and AD skin, the significantly upregulated DEGs in PN included MCEMP1, matrix metalloproteinases (MMPs), OSM, NGF, IL1β, CXCL2, CXCL5, CXCL8, and insulin-like growth factors (IGFB/IGFLs) (Fig. 1C). Significantly downregulated DEGs in PN lesions included CCL13, CCL26, EPHB1, and collagens (COL4/6). Comparing non-lesional skin, PN showed significant upregulation of keratin-family genes (KRT/KRTAP) (Fig. 1D).

**Fig 1.**
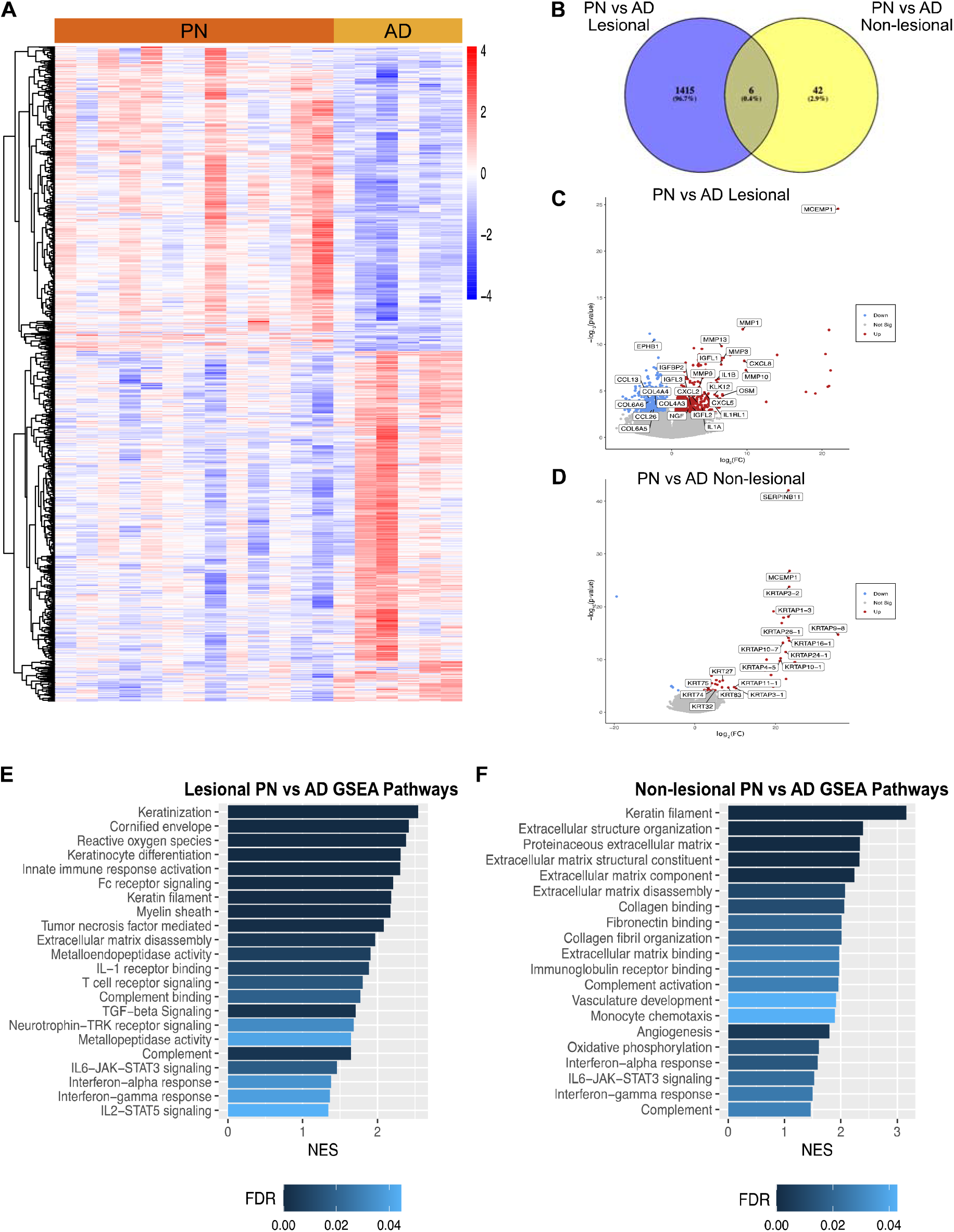
Transcriptomic comparisons of prurigo nodularis (PN) and atopic dermatitis (AD) skin. (A) Heatmap of gene expression for differentially expressed genes (DEGs) between PN lesional vs. AD lesional samples, where red is higher expression and blue is lower expression. (B) Venn diagram of DEGs for PN lesional and AD lesional samples compared to PN non-lesional and AD non-lesional samples. (C) PN lesional vs. AD lesional volcano plot. (D) PN non-lesional vs. AD non-lesional volcano plot. (E) Gene set enrichment analysis (GSEA) for selected significant immune pathways in PN lesional vs. AD lesional skin. (E) GSEA for selected significant immune pathways in PN non-lesional vs. AD non-lesional skin. FC, fold change; FDR, false discovery rate p-values; NES, normalized enrichment score.

GSEA of lesional PN and AD skin revealed that PN lesions had higher enrichment of pathways including keratinization (normalized enrichment score [NES] 2.55, FDR<10^-5^), cornified envelope (NES 2.41, FDR<10^-5^), myelin sheath (NES 2.17, FDR 5.12×10^-4^), TGF-beta signaling (NES 2.09, FDR 0.001), extracellular matrix disassembly (NES 1.97, FDR 0.004), metalloendopeptidase activity (NES 1.90, FDR 0.008), and neutrotrophin-TRK receptor signaling (NES 1.68, FDR 0.033) (Fig. 1E). GSEA of non-lesional PN and AD skin revealed that PN had higher enrichment of pathways including keratin filament (NES 3.16, FDR<10^-5^), extracellular structure organization (NES 3.40, FDR<10^-5^), extracellular matrix disassembly (NES 2.07, FDR 0.01), and angiogenesis (NES 1.99, FDR 0.023) compared to AD (Fig. 1F). These findings of hyperkeratosis in PN compared to AD lesions were corroborated clinically (Fig. 2A) as well as histologically (Fig 2C-F). On immunohistochemical quantification of NGF expression, PN lesional samples had higher dermal NGF than AD lesional samples (6.89% vs 2.51% positive cells, p=.038) (Fig. 2G-H). Furthermore, in non-lesional PN samples, there was greater epidermal compared to dermal NGF expression (16.93% vs 0.87%, p=0.014).

**Fig 2.**
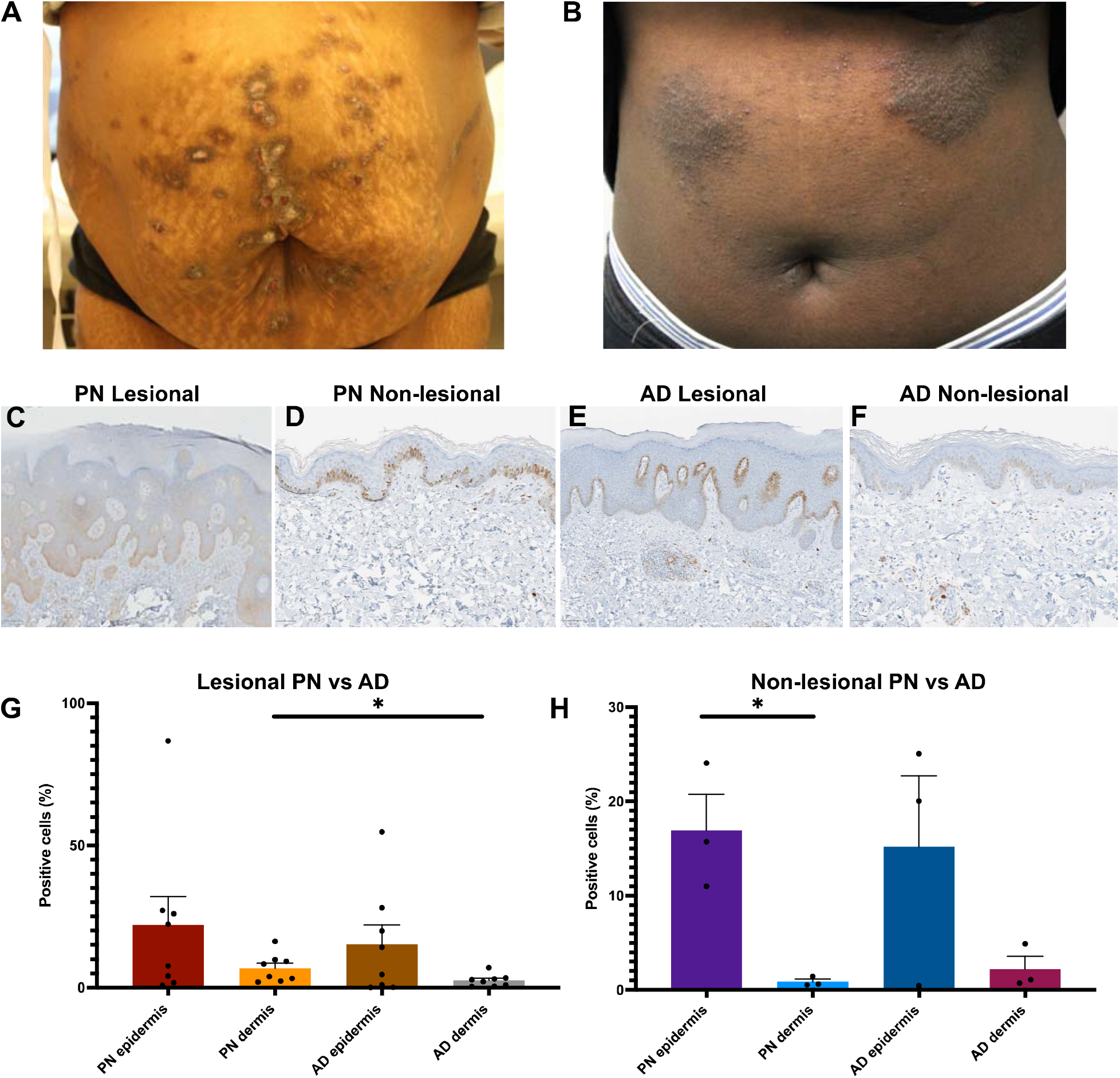
Immunohistochemistry (IHC) analysis of nerve growth factor (NGF) in prurigo nodularis (PN) and atopic dermatitis (AD) skin. (A) Clinical image of a PN patient with fibrotic and hyperkeratotic lesions on the abdomen. (B) Clinical image of an AD patient with lichenified lesions on the abdomen. (C-F) Representative NGF IHC staining in PN lesional, PN non-lesional, AD lesional, and AD non-lesional skin, respectively. Figures C, E: 10x. Figures D, F: 20x. (G) Quantification of the percentage of NGF-positive cells in lesional PN vs. lesional AD skin (n=8 for each condition). (H) Quantification of the percentage of NGF-positive cells in non-lesional PN vs. non-lesional AD skin (n=3 for each condition). *p<0.05.

This study revealed significant enrichment of extracellular matrix remodeling and neurotrophic signatures in PN compared to AD. NGF in the skin is crucial for the survival and regeneration of damaged cutaneous sensory nerves, and transcriptomic and immunohistochemical analysis demonstrated greater NGF expression in PN lesional dermal skin compared to AD. Prior studies have found decreased intraepidermal nerve fiber density in PN skin and increased numbers of NGF-positive papillary dermal nerve fibers compared to controls.^3^ While we found that NGF expression in the PN epidermis is comparable to that in AD epidermis, we additionally found that NGF upregulation is more pronounced in PN lesions than in dermal AD lesions. Transcriptomic analysis also revealed dysregulation in other neurotrophic modulators such as insulin-like growth factors (IGF/IGFL), Il.-I β, and ephrin receptor B1 (EPHB1). These results suggest that PN patients experience greater degrees of cutaneous neural dysregulation compared to AD.

Furthermore, neural dysregulation in PN can be potentiated by alterations in the extracellular matrix. We found that PN lesions had decreases in collagen VI and increases in oncostatin M (OSM), and matrix metalloproteinases (MMPs) compared to AD lesions. Studies have shown that lack of collagen VI, which is necessary for maintaining nerve function and regeneration, can delay peripheral nerve regeneration.^4^ OSM, a cytokine with roles in proliferation or differentiation of hematopoietic and neuronal cells, can also modulate extracellular matrix components and maintain chronic inflammation.^5,6^ These findings are concordant with human clinical trials to date that OSM inhibition has greater efficacy in PN than AD.^7^ OSM can also upregulate MMP activity, which in turn enhances inflammation through degradation of extracellular structures, enabling immune cells to enter and exit skin, and proteolytic activation of cytokines and chemokines.^8–10^ Immunomodulators activated by MMPs include IL-1β, CXCL5, and CXCL8,^8^ whose genes were upregulated in lesional PN skin. Alterations in extracellular matrix components can therefore be major contributors to enhanced inflammation, fibrosis, and neural dysregulation in PN.

In conclusion, we present novel findings demonstrating dysregulation of neural regeneration and extracellular matrix remodeling signatures in PN compared to AD patients. Limitations of this study include patient recruitment from a single tertiary care center, restricting generalizability. Nonetheless, these findings provide deeper insight into the differences in pathogenesis between PN and AD and may aid in the identification of future therapeutic targets.

## Notes

**Funding sources:** SGK is supported by the National Institute of Arthritis and Musculoskeletal and Skin Diseases of the National Institutes of Health under Award Number K23AR077073-01A1. The content is solely the responsibility of the authors and does not necessarily represent the official views of the National Institutes of Health.

**Conflicts of interest:** Shawn G. Kwatra is an advisory board member/consultant for Abbvie, Celldex Therapeutics, Galderma, Incyte Corporation, Pfizer, Regeneron Pharmaceuticals, and Kiniksa Pharmaceuticals and has served as an investigator for Galderma, Kiniksa Pharmaceuticals, Pfizer Inc., and Sanofi.

### Competing Interest Statement

Shawn G. Kwatra is an advisory board member/consultant for Abbvie, Celldex Therapeutics, Galderma, Incyte Corporation, Pfizer, Regeneron Pharmaceuticals, and Kiniksa Pharmaceuticals and has served as an investigator for Galderma, Kiniksa Pharmaceuticals, Pfizer Inc., and Sanofi.

